# Deep Learning of Markov Model Based Machines for Determination of Better Treatment Option Decisions for Infertile Women

**DOI:** 10.1101/606921

**Authors:** Arni S.R. Srinivasa Rao, Michael P. Diamond

## Abstract

In this technical article, we are proposing ideas those we have been developing of how machine learning and deep learning techniques can potentially assist obstetricians / gynecologists in better clinical decision making using infertile women in their treatment options in combination with mathematical modeling in pregnant women as examples.

---

Machine learning approaches in medicine, where applicable, will have tremendous impact on monitoring patients’ health, especially, when care providers want to do the following: improve the precision of the treatment impacts, reduce human errors that could be due to repeated and routine human actions, provide consistency in treatment prescriptions, or any combinations of these. In this commentary, we highlight machine learning (ML) approaches and Artificial Intelligence (AI) based decisions that we believe will have increasing prevalence in clinical care throughout our discipline and others. For the purpose of this discussion, we provide example of infertility treatment for assisting in patients’ healthcare when they present to them obstetrician/gynecologist.

## Infertility Care Options

Suppose we develop an algorithm that can be used in arriving at a decision on which of the treatment options are appropriate for an infertile couple, that can maximize her probability of delivering a child. The data that would be fed into this algorithm usually consists of clinical, demographic and genetic level information from an individual infertile couple for whom a decision on treatment options is to be provided by algorithm. This information we call patients’ level data.

Machines using these algorithms are equipped with parameters whose values were computed with prior information on large population level historical infertile couples’ data who successfully delivered a child with one of the treatment options available, say either IVF or through ovarian stimulation with ovulation induction medicines. There are also types of ovulation induction medicine whose details are fed into the algorithm to be used in the decision making process. These parameters are associated with probabilities of conception, and then for delivery of a baby for various types of infertile couple. One of the methods to obtain such parameters is through Markov Chain based transition probabilities matrices (1–3). Applications of Markov modeling for infertility related data can be found in (4–5) and historically original applications of stochastic processes in biological sciences can be read in (6–8). See Appendix II for the proposed method of computing parameters through Markov Chains approach.

Machines which have the above mentioned algorithms with parameters values obtained from treatment outcomes of a large number of infertile couple we call machine learning algorithm for fertility treatment outcome (MLAFTO). MLAFTO will predict the chances of conceiving and delivering for a new couple who have come to an infertility clinic. MLAFTO would be able to compute probabilities for all the combinations of patient level characteristics combined with their treatment options (See Table 1 for an idea about combination of patient data with treatment options).

Once the chances of predicting the outcomes are computed (see the Appendix for details), then through induction algorithm (i.e. from known values of probabilities of outcomes) the lowest probability of delivering a baby corresponding to a specific treatment option combination, to the highest such probability among all possible sets of probabilities, are sorted out. This procedure through the use of an induction algorithm to suggest a best treatment option we call here as AI based treatment option for an infertile couple.

**Table 1.** Data Used for Computing Probabilitites to Train a Machine

| Patient Characteristics |  |  |  | Treatment Options |  |  |  |  |  | Pregnancy Outcomes |  |
| --- | --- | --- | --- | --- | --- | --- | --- | --- | --- | --- | --- |
| Clinical | Demographic | Genetic | Other | $OI_1$ | $OI_2$ | $OI_3, \dots$ | $OI_p$ | IVF | | Conceived | Live birth |
| $c_1$ | $d_1$ | $g_1$ | $o_1$ | $y/n$ | $y/n$ | $y/n$ | $y/n$ | $y/n$ | | $y/n$ | $y/n$ |
| $c_2$ | $d_2$ | $g_2$ | $o_2$ | $y/n$ | $y/n$ | $y/n$ | $y/n$ | $y/n$ | | $y/n$ | $y/n$ |
| ⋮ | ⋮ | ⋮ | ⋮ | ⋮ | ⋮ | ⋮ | ⋮ | ⋮ | | ⋮ | ⋮ |
| $c_k$ | $d_l$ | $g_m$ | $o_n$ | $y/n$ | $y/n$ | $y/n$ | $y/n$ | $y/n$ | | $y/n$ | $y/n$ |

Note: There are $k$ –clinical, $l$ –demographic, $m$ –genetic, and $n$ –other types of patients characteristics, where the values of $k$ , $l$ , $m$ , and $n$ need not be the same. A given patient could have any one of the all possible combinations ( $k \times l \times m \times n$ ) of patient characteristics, where × is the symbol for multiplication, for example, $\{c_5, d_4, g_1, o_1\}$ , $\{c_k, d_1, g_3, o_7\}$ , $\{c_1, d_3, g_6, o_2\}$ etc., There are $p$ types of ovulation induction (OI) treatments and IVF options available and these are recorded as presence ( $y$ ) or absence ( $n$ ) corresponding to each of the combination of patient characteristics. Finally for each of these patient treatment combinations whether the couple conceived ( $y$ ) or not conceived ( $n$ ) and if conceived then whether a live birth observed ( $y$ ) or not ( $n$ ) is reported.

The Artificial Intelligence quotient of a machine assists in decision making processes for deciding which treatment option(s) are best suited for an infertile woman or a couple who comes to an infertility clinic (based on large historical data and the data on new couple). There are design issues involved on the type of information for the past data which we want to use. That is, whether to use deep learning of the historical data before AI based decision is implemented, or to use machine learning algorithm developed on the past data before AI based decision making can be initiated. AI can be manufactured from deep learning or from machine learning algorithms (Figure 1). In deep learning, each time an infertile couple comes to a clinic, AI will get freshly manufactured from all the data that was fed into the MLAFTO prior to this woman (or couple) and then assists to arrive at a conclusion on which of the treatment options are best suited for the couple. See Appendix III for the key differences between two approaches. That is, the probabilities of conceiving and delivering a baby are computed afresh based on all the infertile couple treatment data (as in Table 1) prior to the new couple who came for infertility care. When we have decided to go with MLAFTO (without deep learning), a fixed set of past data on the infertile woman (or a couple) (as shown in Table 1) will be used to obtain probabilities of conception and delivering a live birth, and then a newly arrived couple’s characteristics will be matched with existing combinations as in Table 1 to suggest a best treatment option for the couple. That is, in the machine learning approach, probabilities of conceiving and for a live birth are computed *a priori* (from a fixed historical data without any further updates), and when a new couple comes for an infertility care, then new couple’s characteristics are matched with existing characteristics of the fixed historical data for which *a priori* probabilities were computed. Through a step by step matching of new couple characteristics with the existing characteristics (inductive reasoning) the closest characteristics that match to the past data is decided. For this closest match the corresponding probabilities computed *a priori* are suggested as possible probabilities for the new couple.

**Figure 1.**
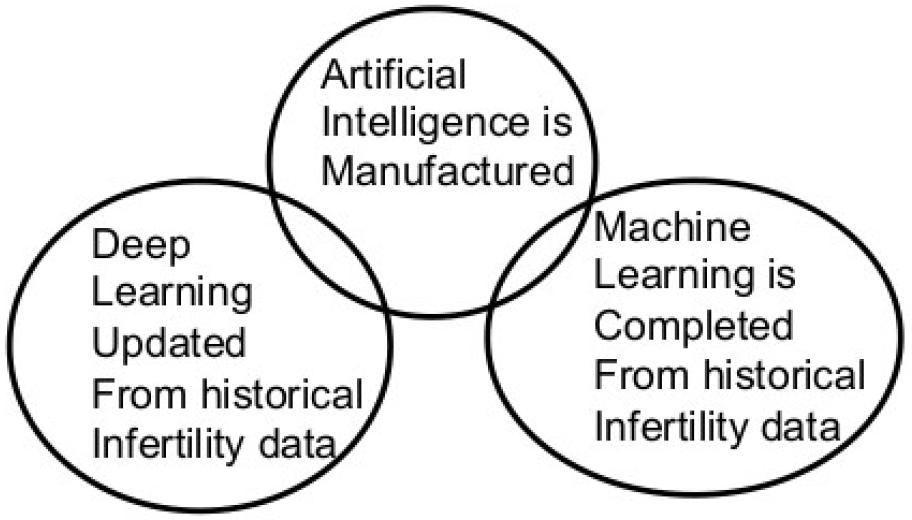
Artificial Intelligence is manufactured from deep learning algorithms or machine learning algorithms or from the both

**Figure 2.**
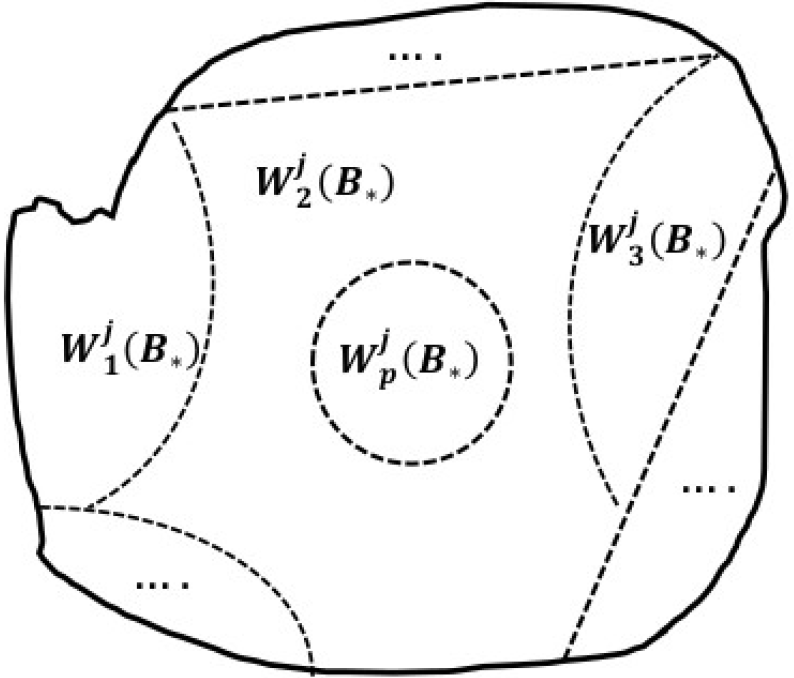
Disjoint sets of infertile women across all the treatment options within the background characteristics(***B***_*_)

Examples of clinical variables include, the duration of infertility, information on earlier treatments if any, depression, anxiety, etc, Potential demographic variables are, namely, age, level of education, duration of relation between couple who came for treatment, etc, genetic variables include: family history of infertility, family history of abortion, miscarriage, etc, other variables include, financial status of the couple, income level of the couple, etc,

Deep learning updates past treatment effects data continuously whereas machine learning computes parameters based on fixed data. However, machine learning parameters can be also updated as newer data becomes available (and also should there be a need to update parameters those were already computed due to a change in treatment options or for some other reason).

Various information on clinical variables and respective treatment outcomes of the past indicated across the big data sets will need to be continuously updated to construct intelligence on the future treatment options. This continuous updating is done wither through pre-constructed models or through deep learning. Some of the several variables like anxiety, depression, level of education, genetic information could vary across the populations in the world, and simplification of their role in treatment effects could lower the chances of creating precise intelligence and for predicting the capacity of treatments of various populations across the world. We suggest that the information on the variables needed to be shared across various populations for capturing better intelligence. The flexibility of the models can be incorporated through model selection procedures to avoid bias. The advantages of deep learning will be maximized with the usage of big data sets and the conception and baby delivery probabilities can be precisely predicted. A word of caution is that deep learning on big data would not automatically guarantee to gather precise AI for predicting the treatment effects and these big data have to be structured accurately with mathematical constructions to be able to build models.

## Acknowledgements

We thank the following individuals in the alphabetical order of their last name for very valuable comments: Medina Jackson-Browne (Brown University, Providence), N.V. Joshi (Indian Institute of Science, Bangalore), K. Praveen (Microsoft, Irvine), P. Sashank (CEO, Exactco, Hyderabad). Actual journal submission contain more examples and results.

## Appendix I: Machine Learning Algorithm for Fertility Treatment Outcome(MLAFTO)

Let *x*({*c*_*_, *d*_*_,*g*_*_, *o*_*_}) be the probability that an infertile woman with characteristics ∗ such that *= 1,2,…,*k* × *l* × *m* × *n* with *i*^*th*^ –ovulation induction (OI) treatment or IVF for *i*= 1,2,…,*p* is conceived or delivered.

We compute *x* values for all * combinations. We compute 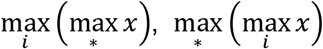. Once the maximum probabilities are computed and ranking of these probabilities over various ∗ and *i* will provide relative chances of conceiving and delivering a live birth. AI quotient will match these combinations of ∗ and *i* is with the new couple who come to the clinic and suggest the probability of conceiving and delivering a live birth. See Appendix II for computing probabilities and descriptions related to *max* functions.

## Appendix II: Computation of Probabilities Through Markov Chains

In this Appendix, we propose a Markov Chain based approach in computing probabilities of conception and delivering a live birth under various treatment options.

Suppose we want to compute the probability of conception and then delivering a baby for an infertile woman with a combination of background variables, say, {*c*_5_, *d*_4_, *g*_1_, *o*_1_} and with *OI*_*i*_ *or IVF* treatments explained in the paper. Let *B*_1_ be the set of all infertile women with background variables B_1_={{*c*_5_, *d*_4_, *g*_1_, *o*_1_} who will be *OI*_*i*_ *or IVF* treatment options. Let 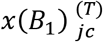 be the probability that an infertile woman at the state *j* with characteristics {*c*_5_, *d*_4_, *g*_1_, *o*_1_} and with *i*^*th*^ – ovulation induction (OI) treatment or *IVF* for *i* = 1,2,…,*p* is conceived in *T* – time steps. Let 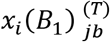 be such a probability to deliver a baby, and *x*(*B*_1_)_cb_ be the probability of baby is born to a woman within *B*_1_ given that the woman is conceived. These three probabilities can be computed using below formulas:

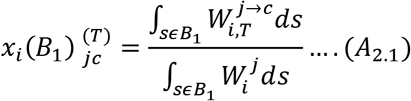

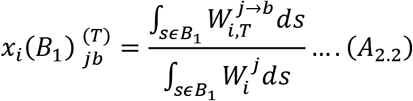

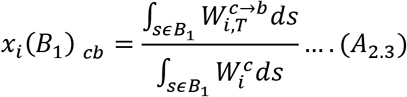

where 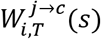 denotes *s*^*th*^ infertile woman in the state *j* who is on i^th^ treatment conceives in *T* – time steps, 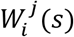 denotes *s*^*th*^ infertile woman in the state *j* who is on i^th^ treatment. 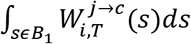 is the total number of infertile women in the set *B*_1_ who have moved from the state *j* to the state *c* who are on i^th^ treatment and 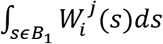 is the total number of women in the set *B*_1_ who are at the state *j* who are on i^th^ treatment. Suppose *B*_2_ be another set of all infertile women with different background variables, say, *B*_2_={{{*c*_1_, *d*_3_, *g*_6_, *o*_2_} with *OI*_*i*_ *or IVF* treatments}, then we can compute corresponding transition probabilities by a similar type of formulas as in (A_2.1_) − (A_2.2_).

Probability of transition from the state *c* to the state *b* does not depend upon 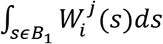 but only on the 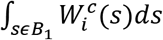, so the random variable responsible for the transition between these two states, say *Y*, obeys Markov property. Moreover, the transition probability matrix *P*_*i*_(*B*_1_) for the set of infertile women *B*_1_ between states {*j*,*c*,*b*} who are on i^th^ treatment can be written as,

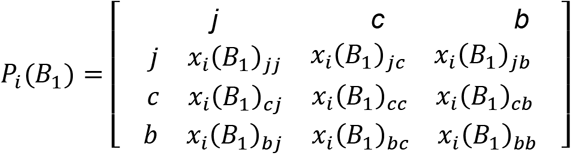

where *x*_*i*_(*B*_1_)_*jj*_ + *x*_*i*_(*B*_1_)_*jc*_ = 1, *x*_*i*_(*B*_1_)_*cc*_ + *x*_*i*_(*B*_1_)_*cb*_ = 1 and *x*_*i*_(*B*_1_)_*bb*_ = 1. *x*_*i*_(*B*_1_)_*jb*_ = 0 due to Markov property. Where as *x*_*i*_(*B*_1_)_*cj*_ = *x*_*i*_(*B*_1_)_*bj*_ = *x*_*i*_(*B*_1_)_*bc*_ = 0 due to transition from *c* → *j*, *b* → *j*and *b* → *c* are impossible. Similarly, we will compute

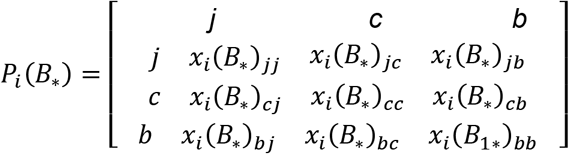

for *= 1,2,…,*k*×*l*×*m*×*n*. Let 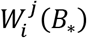 be the number of infertile women (state *j*) within background characteristics *B*_*_ who are on i^th^ treatment and let *W*^*j*^(*B*_*_) be the total number of infertile women within background characteristics *B*_*_ such that

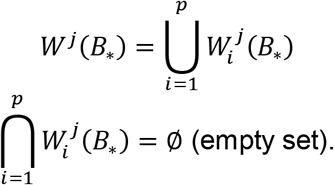

Once *P*_*i*_(*B*_*_) is computed based on certain design of the sample population, the sizes of 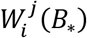 are not changed for computing probabilities using (A_2.1_) – (A_2.2_). That is, the matrix *P*_*i*_(*B*_*_) is not updated based on newer women who have started treatment after the designed time interval

## Two functions

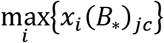 and 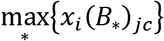

The function 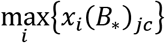 describes that the maximum of the probability values of women with background characteristics *B*_*_ across all the treatments, which is obtained as

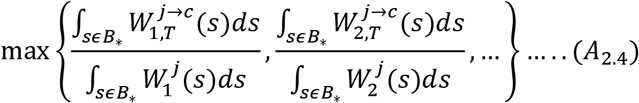

Through the expression (A_2.4_) we will obtain *k* × *l* × *m* × *n* maximum values, where each maximum value represents maximum probability of conceiving by an infertile woman from a particular set of background characteristics and corresponding treatment type for which this maximum value is obtained. Similarly, we can construct 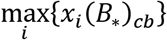. The function 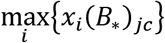 describes that the maximum probability of conceiving within the women who are i^th^ treatment across different background characteristics, which is obtained as

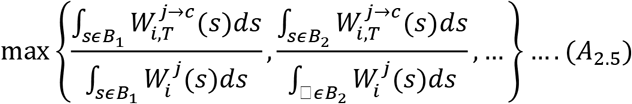

Let 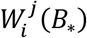 be the number of infertile women (state *j*) within background characteristics *B*_*_ who are on i^th^ treatment and let *W*^*j*^(*B*_*_) be the total number of infertile women within background characteristics *B*_*_, then

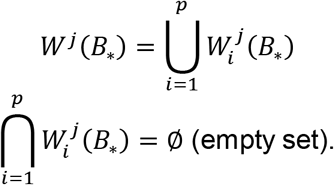

See also Figure 2 to see this disjoint property of infertile women within each background characteristics.

## Result

Total infertile women with background characteristics {*B*_*_} can be written as the union of disjoint sets of women across all treatment options, i.e.

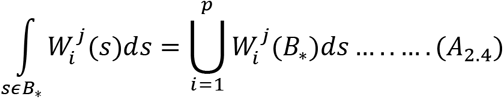

## Appendix III: Machine Learning Versus Deep Learning in Computing Probabilities of Conception and Delivery

Suppose a new infertile woman whose background characteristics {*B*_*N*_} is interested to start one of the available treatments *OI*_*i*_ *or IVF*. Let us understand how machine learning techniques are applied to decide which of the treatment will give maximum chance of conception and delivering a baby. Prior to a decision making process on treatment options for this woman, let us suppose that probabilities of conception and delivery were previously computed through MLAFTO explained in the Appendix I and *P*(*B*_*_) for all * in the Appendix II. The data used for these two computations is usually a predetermined or pre-designed one, i.e. the time frame and other design aspects of the data were well defined and are without any data related errors. MLAFTO matches the new infertile woman characteristic set *B*_N_ with the sets {*B*_*_ : *= 1,2,…,*k* × *l* × *m* × *n*}. Let {*B*_y_} be the set that matches with the new woman characteristics such that {*B*_y_} − {*B*_N_} = ∅ (null set). The corresponding values of

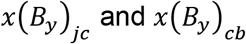

are considered as chances of conception and chances of delivery for the new woman who came to the clinic.

Note that, the success or failure data of woman with {*B*_N_} is not used in computation of *P*(*B*_*_) for all * which is the key for machine learning type of algorithm.

If each treatment trial of a woman whether or not that woman conceives is considered as one time step of treatment (or 1-cycle of treatment) and the duration from conceiving of a woman to whether or not a baby is delivered is considered as one time step of pregnancy (or 1-cycle of pregnancy), and let 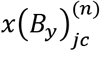 and 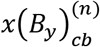 be the corresponding *n* – step or *n* –cycle probabilities, then by Markov property, we have

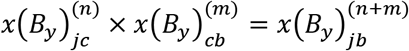

When another infertile woman with background characteristics {*B*_M_} comes to the clinic for the purpose of decision making of which type of treatment will be needed for a successful delivery, the prior computed transition probability matrix *P*(*B*_*_) for all * that was used in matching for a woman with {*B*_N_} was not updated with the success or failure information of the woman with {*B*_N_}. In a way, the matrices *P*_*i*_(*B*_*_) are static in case we are using machine learning algorithms and these are not influenced by new data generated on newer infertile women who come to the clinic after constructing *P*_*i*_(*B*_*_).

Once an infertile woman walks into the clinic with background characteristics {*B*_M_}, if deep learning techniques are implemented to predict the probabilities of conceiving (say, *y*(*B*_N_)_jc_) and the delivery (say, *y*(*B*_N_)_jc_), then the computations of such probabilities are different than machine learning techniques. Each time a new infertile woman with {*B*_M_} comes to the clinic for the treatment purposes, instead of matching procedure with the existing static model explained above, deep learning involves reconstructing of the transition probability matrices *P*_*i*_(*B*_*_), for *i*=1,2,…,*p* for conceiving and delivery with whatever data that is available prior to arriving of the woman with {*B*_M_}. Rest of the computational procedures explained in the Appendix II remains the same. Deep learning techniques usually delay the output due to reconstructing of the *P*_*i*_(*B*_*_ each and every time a new infertile woman comes to clinic.

General introductions of machine learning techniques, motivations, and key ideologies were explained in a variety of research areas can be found in (9–13). Specific ideas related to deep learning techniques were also well developed (14), deep learning techniques and applications were summarized (15) and an overview of importance of machine learning algorithms in medicine can be found in (16). As explained in our article, the machine learning and deep learning techniques broadly use the same data within the specific goals, but their approach of handling the data and models distinguish them from each other. Statistical thinking had contributed several aspects of machine learning, for example, in developing computationally intense data classification algorithms, methods in data search and matching probabilities, data mining techniques, model classification and model fitting algorithms, and a combination of all these (see for example, (17–29) and for a collection of articles related to statistical methods in machine learning see (30). Model-based machine learning methods (31) and the construction of coefficients in a regression model can be benefited by machine learning methods (32).

Deep learning techniques, instead of focusing on model-based approaches, they would assist in understanding intricate structures of the large data sets and various interlinkages between these data sets (33). Importance of unsupervised pre-training to the structural architecture and the hypothesis of testing design effects of such experiments are well studied (34, 35). Deep learning and machine learning techniques could also assist in questions related to health informatics, disease detection, item response theories, and bioinformatics research (36–41). There were also successful methods in deep learning algorithms which score patients in ICU (Intensive Care Unit) for their severity and to predict mortality without using any model based assumptions in scoring systems (42) and for other medical applications, for example detection of worms through endoscopy (43), ophthalmology studies (44), cardiovascular studies (45), Parkinson‘s disease data (46), medical scoring systems (47). Deep learning procedures involved in various levels of abstraction for ranking system models can be found in (48, 49), applications for mathematical models, parameter computations and stability of algorithms are found in (50–56).

Statistical and stochastic modeling principles were applied in deep learning algorithms to strengthen the object search capabilities or for improved model fitting in uncertainty(57, 59). Boltzmann machines assist in the deep understanding of the data by linking layer level structured data and then by estimating model parameters through maximum likelihood methods (60, 61). Random backpropagation and backpropagation methods help in stochastics transition matrix formations and computing quicker search algorithms in higher dimensional stochastic matrices and literature related to backpropagation could be found in several places, for example, see in (62–65). A survey of statistical learning algorithms and their performance evaluations can be found in (66).

## Appendix V: Theorems

### Theorem A.1

When *W*^*j*^ is the total number of infertile women (state *j*) whose data is used in the machine learning algorithm and *δ* ∈ [1,*klmn*] and *α* ∈ [1,*p*] are considered as continuous for background characteristics and treatment options, then

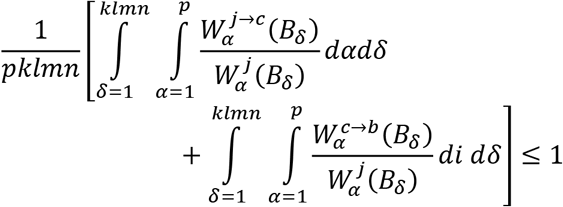

***Proof***: We have,

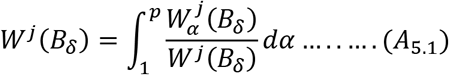

and

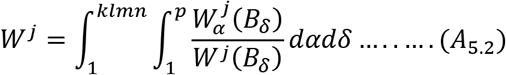

Note that,

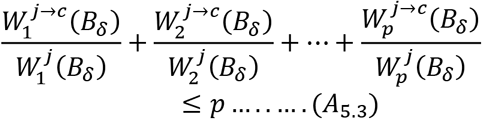

and

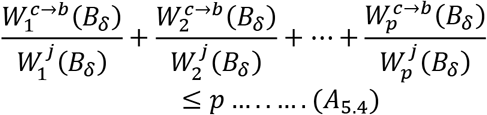

from the inequality (*A*_4.3_) we can obtain,

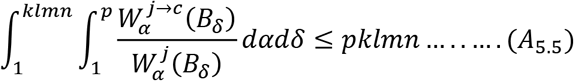

from the inequality (*A*_4.4_) we can obtain,

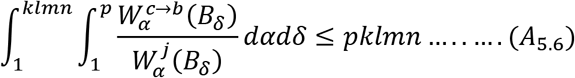

Required result is deduced from two inequalities (*A*_4.5_) and (*A*_4.6_).

### Theorem A.2

For continuous *α* and *δ*, we have

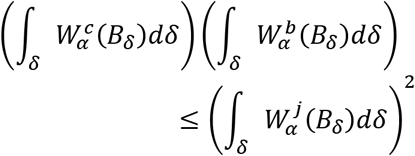

***Proof***: We know,

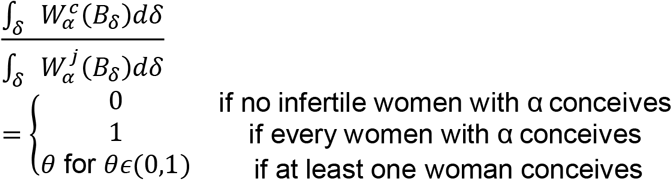

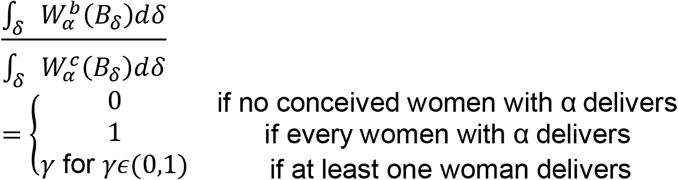

These imply,

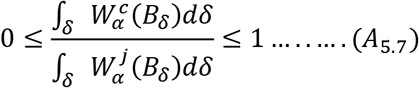

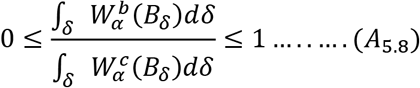

From (A_5.7_) and (A_5.8_), we can deduce required result.

### Theorem A.3

Let *f*:*A* → ℝ^+^ and *g*: *B* → ℝ^+^ where *A* is the set of fractions of (A_5.7_) and *B* is the set of all fractions of (A_5.8_), then *f* and *g* are defined only at the adherent points of *A* and *B*, respectively.

***Proof***: Note that,

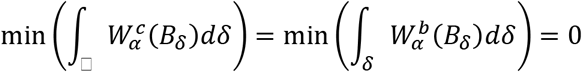

and

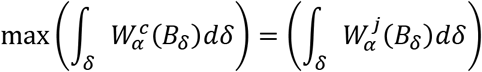

Two sets *A* and *B* are constructed from (A_5.7_) and (A_5.8_) as

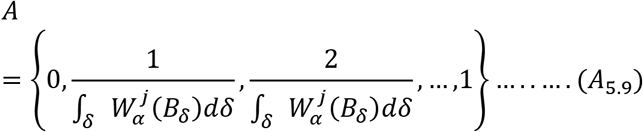

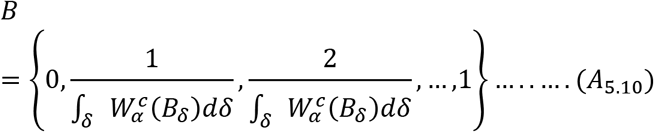

From the elements of the set *A* as in (A_5.9_), *f* is not defined at open subintervals

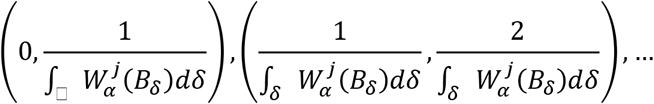

and from the elements of the set *B* as in (A_5.10_), *g* is not defined at open subintervals

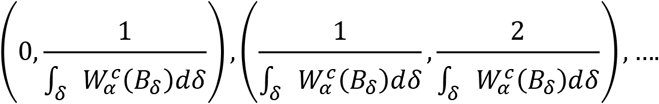

Hence, *f* and *g* are defined only at the adherent points of *A* and *i*

